# Photoperiodic history modulates the response of the saccus vasculosus transcriptome to seawater exposure in Atlantic salmon

**DOI:** 10.1101/2025.03.14.643326

**Authors:** Alexander C West, Shona Wood, Marianne Iversen, Jayme van Dalum, Even H Jørgensen, Simen R Sandve, David G Hazlerigg

## Abstract

Atlantic salmon (*Salmo salar*) move from fresh- to seawater environments following a seasonally timed preparative transition called smoltification, which takes place under photoperiodic control in the freshwater environment. In masu salmon (*Oncorhynchus masou*), coordination of photoperiodic sexual maturation is proposed to involve in a fish-specific circumventricular organ, the saccus vasculosus (SV), through its intrinsic opsin-based light sensitivity, thyrotrophin secretion and modulation of deiodinase activity (TSH-DIO cascade). The saccus vasculosus is a highly vascularized structure located on the ventral side of the hypothalamus and its interface between the blood and cerebrospinal fluid also hints at a role in ionic balance of the cerebrospinal fluid (CSF). Both the potential photoperiodic and ionic functions of the SV led us to perform transcriptome analysis of the SV in smoltification in Atlantic salmon. Our data show that SV response to seawater exposure is highly dependent on photoperiodic history and identifies ependymin as a major secretory output of the SV, consistent with a role in control of CSF composition. Conversely, we could not detect crucial elements of the opsin-TSH-DIO cascade suggesting that the photoperiodic history-dependence of the SV to seawater exposure is unlikely to stem from SV-intrinsic responses to photoperiod.

## 1. Introduction

Annual changes in day-length (photoperiod) alter the seasonal environment, particularly at higher latitudes. As a result, many organisms compartmentalize elements of their life history to specific times of year ^1^. Atlantic salmon (*Salmo salar*) undergo two major seasonally timed developmental processes in their lifecycle: reproduction and smoltification ^2,3^. Reproduction occurs following migration to their natal streams in the autumn ^2^. Smoltification occurs earlier in life when immature ‘parr’ salmon prepare to leave their freshwater environments and migrate to the ocean during the spring. While smoltification is a complex event that includes changes in physical appearance, growth, endocrinology and immune function ^4–6^, its most striking feature is the change in osmoregulatory function which allows smolts to thrive under the osmotic gradient of their seawater environment ^7^.

Atlantic salmon, like other vertebrates, time seasonal changes in phenotype by integrating photoperiod information ^1^. Short photoperiods (eg. 8L:16D) provide a winter signal and long photoperiods (eg 18L:6D; or constant light, LL) provide a summer signal. Current working models for the physiological mechanism underlying the vertebrate physiological response focus on thyrotrophin signaling and hypothalamic changes in thyroid hormone (TH) metabolism ^8–10^. In photoperiodic birds and mammals, decoding of photoperiod takes place in the pars tuberalis (PT), a specialized pituitary tissue^11^. On sensing summer-like daylengths, the PT rapidly produces a modified form of thyrotrophin (TSH) whose retrograde action in hypothalamus stimulates the expression of deiodinase type II (*Dio2*), a thyroid hormone activating enzyme, within specialized tanycyte cells lining the third ventricle. The induction of DIO2 and concomitant suppression of DIO3, a thyroid hormone inactivating enzyme, leads to the conversion of the biologically inert T4 form of thyroid hormone to the bioactive variant T3. Local T3 elevation within the hypothalamus then orchestrates a neuroendocrine program that stimulates organism-specific, summer-type physiological states. In contrast, short winter-like photoperiods lead to low levels of PT-type TSH and DIO2, and increased DIO3 levels reduce the hypothalamic T3 concentration which is linked to winter-type physiological states ^12–15^. While teleosts do not have an anatomically distinct PT ^16^, two models nonetheless place seasonally dependent changes in TSH expression at the core of the photoperiod integration mechanism. Based on studies in masu salmon, Yoshimura and colleagues have described the presence of components of the TSH-DIO pathway, including light-sensitive opsins, in coronet cells of the saccus vasculosus (SV), a fish-specific circumventricular organ (**Figure 1A-C**) ^17^. In addition, they reported that surgical removal of the SV in masu salmon (*Oncorhynchus masou*) blocks winter photoperiod-induced testicular growth, suggesting that the SV may be a seasonal sensor of photoperiod and coordinator of seasonal reproduction in salmonids ^17^. Contrastingly, in Atlantic salmon, in has been reported that a novel TSHβ subunit isoform, designated TSHβb is expressed in the dorsal region of the pars distalis, with levels increasing markedly during photoperiod-dependent smoltification, leading to the proposal that this isoform plays an analogous role to PT-derived TSH in birds and mammals ^18,19^.

**Figure 1.**
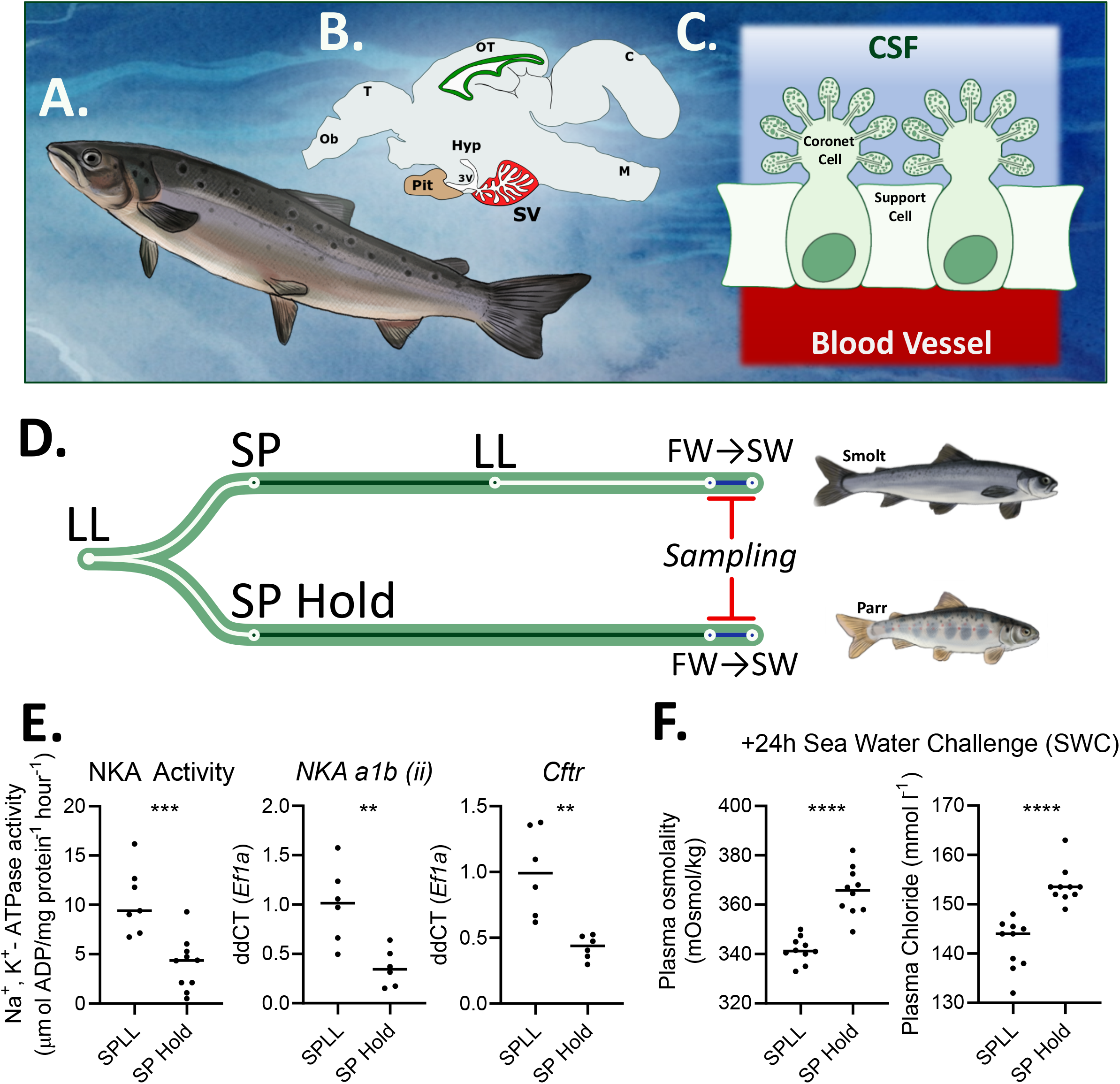
Experimental design and evidence of hypoosmotic potential in SPLL group. (A-C) Atlantic salmon have a well-developed SV located next to the ventral side of the brain, caudal to the pituitary gland and connected to the third ventricle. The SV contains coronet cells whose protrusions into the CSF are covered in several globule structures. Coronet cells are flanked by supporting cells. (D) Experimental design: the SPLL group (top) were transferred to short photoperiods (SP) for six weeks and returned to constant light (LL) for six weeks; the SP-hold group (bottom) were kept on SP for 12 weeks. Both groups sampled in freshwater (FW) and 24h in seawater. (E) osmoregulatory potential of the gill in SPLL and SP hold groups as shown by Na^+^ K^+^ ATPase activity, *NKAa1b(ii)* expression and *Cftr* expression. (F) increased hypoosmoregulatory ability of the SPLL group as shown by plasma osmolality and plasma chloride levels following 24h is seawater. Data in 1E and 1F are replotted from Iversen et al (2020)^65^. Abbreviations: Ob. Olfactory bulb, T. Telencephalon, OT. Optic tectum, C. cerebellum, M. mesencephanlon, Pit. Pituitary, 3V. third ventricle, SV. Saccus vasculosus, CSF. cerebrospinal fluid.

To date, no studies have investigated the relationship between the SV and smoltification. It is notable, however, that the SV coronet cell globules of several teleosts, including rainbow trout (*Oncorhynchus mykiss*), contain high densities of Ca^2+^ containing vesicles which are suggested to act as reservoirs that release calcium ions into the cerebrospinal fluid (CSF) of the third ventricle ^20–23^. The SV is also extensively innervated by neurons that project both to and from the hypothalamus and neurosecretory CSF contacting dopaminergic neurons ^24,25^. While there are no categorical divisions between teleosts from different ecological niches, there is a notable loss or reduction of the SV in most primary freshwater fish contrasting with marine teleosts who usually have well-developed SVs ^26^. The SV of some euryhaline fish changes with seawater exposure. For example, the prevalence of glycosaminoglycans, polysaccharides important for ECM function ^27^, on the cell surface of stickleback (*Gasterosteus aculeatus*) and rainbow trout SV coronet cells is greater when fish are kept on freshwater compared to seawater ^28,29^. Interestingly, injection of SV homogenate from seawater-acclimated rainbow trout into the third ventricle of freshwater-acclimated rainbow trout increased their seawater survival time compared to uninjected animals, supporting a role of the SV in SW adaptation ^30^. Together, these data suggest a role for the SV in CSF ion composition which may be adaptive during marine migration of salmon during smoltification.

To determine if the SV plays a role in photoperiodic control of smoltification and / or osmoregulation, we have conducted an analysis of transcriptomic changes in the SV of Atlantic salmon subjected to a standard photoperiod protocol known to induce smoltification under laboratory conditions. Our data support earlier reports suggesting a role for the SV in calcium homeostasis and indicate that the SV is sensitive to both photoperiod and salinity, with a significant interaction between these factors. Despite this, key components of the TSH-DIO pathway are not expressed in the SV of juvenile Atlantic salmon, implying that these observed changes are secondary to neuroendocrine regulation mediated by other neuroanatomical sites.

## 2. Materials and Methods

### 2.1 Animal Welfare Statement

All studies were performed in accordance with Norwegian and European legislation on animal research. The smoltification experiment was conducted as part of ongoing smolt production at Havbruksstasjonen i Tromsø, which is approved by the Norwegian Animal Research Authority (NARA) for the containment and experiments on fresh and seawater fish. Formal approval of our experimental protocol was not required because the experimental conditions were consistent with routine animal husbandry practices and therefore there was no compromise of animal welfare.

### 2.2 Experimental Design

Atlantic salmon (Salmo salar, Aquagene commercial stain) were raised from hatching in freshwater under continuous light (> 200 lux at water surface) at ambient temperature (∼10°C). Juvenile fish were housed in 500L tanks and fed continuously with pelleted salmon feed (Skretting, Stavanger, Norway). At 11 months of age, 1400 parr (mean weight 40.3g) were distributed among distributed among eight 300 L circular tanks and allowed to acclimate for one week.

Following acclimation, the fish were treated in two groups. Group 1 (SP-hold) were transferred to short photoperiod (SP; 8h light / 24h) for 16 weeks. Group 2 (SPLL) were transferred to SP for 8 weeks before transfer to constant light (LL) for an 8 weeks. Fish were sampled in FW prior to SW transfer, then the remaining fish were transferred to a full-strength seawater tank (34 ‰ salinity) under constant light. After 24h in seawater the final group of fish were collected (see **Figure 1C** for experimental design). The experiments were performed in duplicate tanks, five fish were taken from each duplicate tank at each collection point, sex was not determined.

### 2.3 Tissue collection

On collection, the fish were netted out and lethally anesthetized in benzocaine (150 ppm). Once fish opercula stopped moving, blood was sampled from the caudal vein, then the fish were decapitated and SVs rapidly dissected and stored in RNAlater (Sigma-Aldrich, USA) at 4°C for 24 h, then frozen at −80°C until further processing.

### 2.3 RNA extraction

SVs were selected from two fish per sampling point from duplicate tanks. SV tissue was disrupted using a TissueLyser II (QIAgen, Germany) and RNA extracted using a RNeasy micro kit (QIAgen, Germany). RNA concentrations were measured using a NanoDrop ND2000c spectrophotometer (NanoDrop Technologies, USA) and stored at −80°C until further processing

### 2.4 RNA sequencing

Sequencing libraries were generated using TruSeq Stranded mRNA HS kit (Illumina). Library concentration was measured using a QUBIT BR kit (Thermo Scientific) and mean length was measured using a 2100 Bioanalyzer and DNA 1000 kit (Agilent Technologies). Individual samples were then barcoded using Illumina HiSeq indexes and single-end 100bp sequencing of the samples were performed using an Illumina HiSeq 2500 at the Norwegian Sequencing Centre (University of Oslo, Norway). Cutadapt was next used to remove sequencing adapters, trim low quality bases and remove short sequencing reads under the parameters -q 20 -O 8—minimum-length 40 ^31^. FastQC was used to perform quality control of the reads. Transcripts from single-end RNAseq data were quantified based on the Atlantic salmon reference transcriptome version 3 (Ssal_v3.1, GCA_905237065.2) by using Salmon (v 1.1.0.) on mapping-mode using the flags --keepDuplicates for indexing. The flags –validateMappings, --gcBias and –numBootstraps 100 were used during quantification ^32^. Transcript level counts were uploaded to EdgeR (3.42.4) using the command catchSalmon ^33^. Differential gene expression was then calculated by ANOVA-like contrasts. Three samples were excluded from the analysis due to poor data quality. The final replicate numbers in each group were therefore: SP-hold_FW = 4, SP-hold_SW = 3, SPLL_FW = 2, SPLL_SW = 4. Data is accessible online (GEO accession number GSE291612).

### 2.5 Gene Ontology Analysis and transcription factor binding site analysis

Mouse and zebrafish homologues to Atlantic salmon genes were identified using BioMart ^34^. The target and background sets of genes were then submitted to the GOrilla gene ontology tool along to identify enriched GO terms ^35^. Enrichment for transcription factor binding sites was determined using SalmotifDB ^36^.

## 3. Results

### 3.1 Photoperiodic stimulation of smoltification

To confirm smolt status, we collected gill samples for NKA activity assays and qPCR to infer the preparation of the fish to the osmoregulatory challenges of marine migration. Gill NKA activity was increased in fish collected in the SPLL FW group compared to the SP-hold FW group, typical of fish adopting smolt physiology (**Figure 1E**). In the gill we also observed an increase in mRNA expression of two smolt-associated ion channel subunits: *NKAa1b(ii)* and *Cftr* (**Figure 1E**) ^37,38^. We next measured the ability of the fish to hypo-osmoregulate by measuring of blood plasma osmolality and chloride concentrations following 24h in full strength seawater. Both blood osmolality and chloride levels were increased in the SP-hold group relative to the SPLL group, indicating a reduced capacity of the SP-hold group to defend against the osmotic gradient experienced in seawater (**Figure 1F**). Together these data show that, consistent with previous work, the photoperiod treatments of our experiment were sufficient to stimulate smoltification in the SPLL group but not in the SP-hold group.^39^

### 3.2 Transcriptomic analysis of saccus vasculosus highlights role in ependymin production and exocytosis

We next wanted to examine our RNAseq data for insights into the SV biology of Atlantic salmon. Our SV data identified 2743 transcripts expressed with >30 transcripts per million reads (TPM; **Supplementary Table 1**). We analyzed this gene set to identify over-representation of genes associated with specific gene ontology (GO) terms (**Supplementary Table 1**). The most significant over-represented GO terms were associated with translation (eg. GO:0006412 translation, GO:0003735 structural constituent of ribosome, GO:0005840 ribosome), neurons (eg GO:0045202 synapse, GO:0097458 neuron part) and oxygen transport (eg GO:0031721 hemoglobin alpha binding, GO:0005344 oxygen carrier activity).

We also examined highly abundant transcripts with the expectation that their protein products may have a major role in SV function. Ranking the genes by average TPM highlighted *Ependymin-1* (*Epd1*) and *Ependymin-2* (*Epd2*), calcium-binding extracellular matrix (ECM) proteins^40^, whose combined transcript abundance represented an impressive 6% of the total counts. Other abundant transcripts included: *parvalbumin* (*Prvt*), an intracellular calcium buffer and known SV marker gene ^41,42^; *2-peptidylprolyl isomerase A* (*Ppia*, aka. *Cyclophilin A*) an enzyme which facilitates protein folding and whose secretion is linked to inflammation ^43,44^; and the oxygen transporter *Hemoglobin* (*Hbb*), whose prevalence is likely due to the high vascularization, and therefore increased red blood cell complement, of our SV sample ^45^. We note that there are several highly expressed genes in our dataset whose function remains uncharacterised (**Supplementary Table 1**).

### 3.3 Photoperiod changes the SV transcriptional response to seawater transfer

An ANOVA-like test on SV transcriptomes from SP hold FW, SP hold SW, SPLL FW and SPLL SW groups identified a list of 335 transcripts that were differentially regulated with an FDR cutoff of <0.01 (**Supplementary tables 2-6**). Hierarchical clustering of the groups separated their expression into five major clusters with distinctive expression profiles (**Figure 2**).

**Figure 2.**
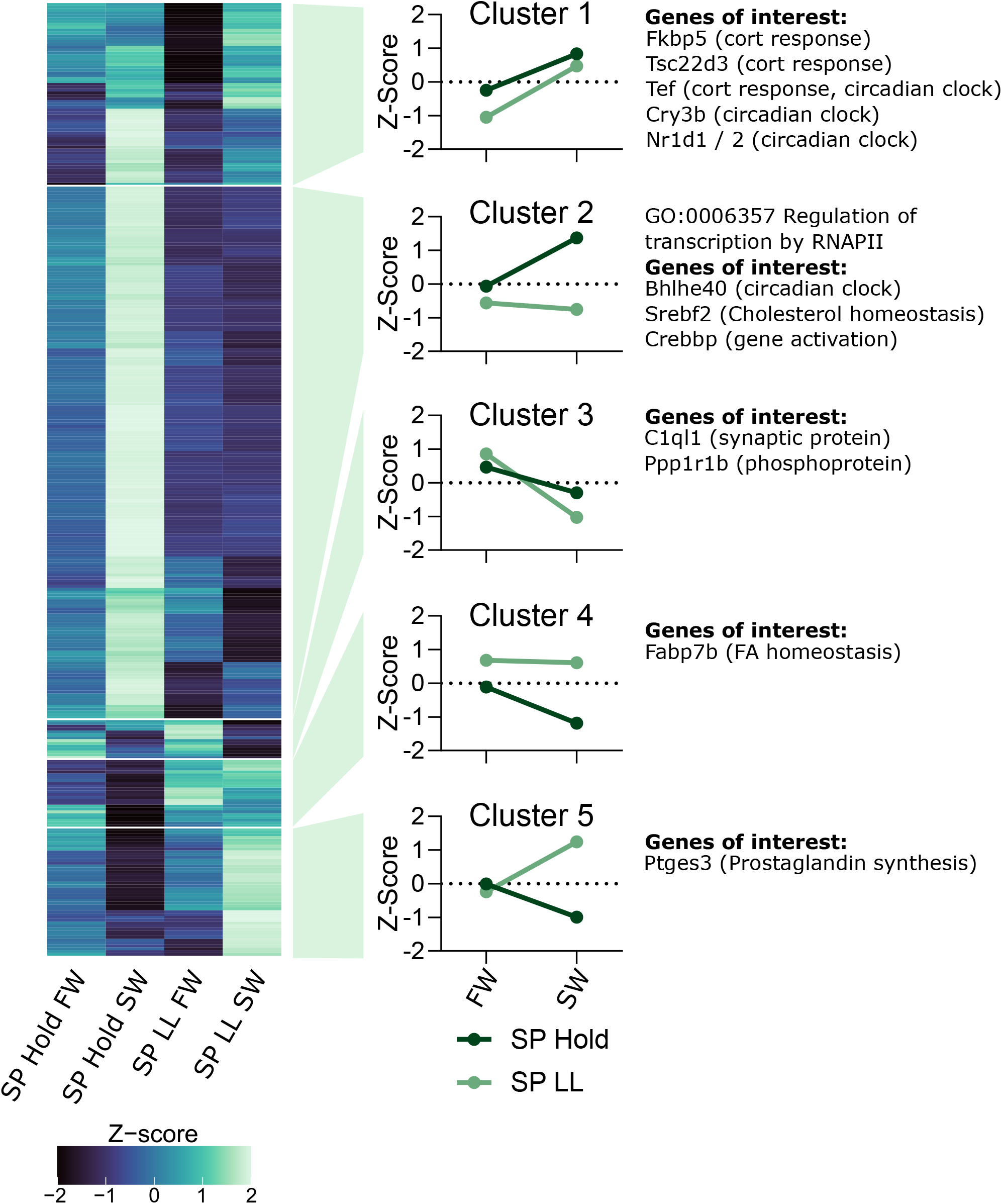
The Atlantic salmon transcriptome is sensitive to both photoperiod and water quality. Heatmap showing the mean expression of differentially expressed genes clustered into five major clusters of gene expression profiles. Specific genes of interest are highlighted next to each clustering group.

Cluster 1 and 3 transcript expression was affected by seawater transfer independent of photoperiod (**Figure 2**). Cluster 1 transcripts were induced by seawater transfer (64 transcripts; **Figure 2**). While there was no significant enrichment of GO terms (FDR cutoff <0.01) or TFBSs (q-value cutoff <0.01), the glucocorticoid receptor binding site was enriched (q-value = 0.055) and the group contained several genes with strong links to cortisol response pathways (eg. Fkbp5^46^, Tsc22d3^47^), some of which are also related to circadian clock function (eg. Tef ^48^). Cluster 3 transcripts, in contrast to cluster 1, were suppressed by seawater transfer (14 transcripts; **Figure 2**). This cluster includes *C1ql1*, a secreted synaptic protein^49^, and *Ppp1r1b*, which encodes *Protein phosphatase 1 regulatory subunit 1B*, also known as *dopamine-and cAMP-regulated neuronal phosphoprotein 32* (*DARPP-32*), which is involved in dopamine and glutamate-dependent synaptic plasticity^50^.

Cluster 4 was increased under SPLL compared to SP hold and comparatively stable following seawater transfer under both photoperiod treatments (24 transcripts; **Figure 2**). The group had no enriched GO terms (FDR cutoff <0.01), but we identified a significant enrichment of the Gmeb1 TFBSs, a glucocorticoid modulatory element^51^. We note, however, that the expression of *Gmeb1* is low (19847^th^ most expressed transcript, average TPM 3.3; **supplementary table 1**).

Cluster 2 and 5, which together make up over two thirds of the differentially regulated transcripts, showed photoperiod-dependence in their response to SW challenge. Cluster 2 transcripts were induced following seawater transfer in the SP hold group, but not in the SPLL group (187 transcripts; **Figure 2**). The group was enriched for several GO terms relating to transcriptional regulation (FDR cutoff <0.01; **Supplementary table 3**; **Figure 2**), and was enriched for 295 TFBS (q-value cutoff <0.01), including several immediate early genes (eg. EGR1, FOS and JUN) and TEF, a PAR ZIP transcription factor linked to the circadian clock and cortisol response in salmon gill tissue ^48,52,53^. Notably, cluster 2 had high expression of the transcription regulators *Bhlhe40* (aka. *Dec1*), a circadian clock repressor linked to photoperiodic timing in mammals ^54^, *Huwe1*, a central coordinator of ubiquitination^55^, and *Srebf2*, a transcriptional regulator of genes involved in cholesterol biosynthesis^56^. Cluster 5 transcripts were suppressed following seawater transfer in the SP hold group and induced by seawater transfer in the SPLL group (45 transcripts; **Figure 2**). The cluster was not enriched for GO terms (FDR cutoff <0.01; Supplementary table 6) or TFBSs (q-value cutoff <0.01), but did express high levels of *Ptges2*, an enzyme that catalyzes the conversion of prostaglandin to its bioactive E2 form and can influence vascular tone^57^.

### 3.4 The TSH-DIO pathway in the SV is incomplete and not consistently regulated by photoperiod or water salinity

To determine if the TSH-DIO pathway (**Figure 3A**) was present or differentially regulated by photoperiod or water salinity, we extracted counts from each component of the pathway from our transcriptomic dataset. We first checked for the expression of opsins, a family of conserved photosensitive G-protein coupled receptors. Out of the 54 known Atlantic salmon opsins we detected the transcript expression of two: *Opn8a1* and *Rgrb* (**Figure 3B**) ^58,59^. We next checked for the expression of the genes that encode the TSH protein heterodimer and the TSH receptor, TSHR. Atlantic salmon have two *Tshβ* gene duplicates (*Tshβa* and *Tshβb*)^60^ and two alpha subunit gene duplicates (*Glha1* and *Gpha2*), however, none of these four genes or *Tshr* were expressed in our samples (**Figure 3A**). We did, however, detect the expression of *Dio2* (both duplicate genes *Dio2a* and *Dio2b*) and *Dio3* (**Figure 3C**), but for none of these transcripts did we see significant effects of photoperiod or SW exposure.

**Figure 3.**
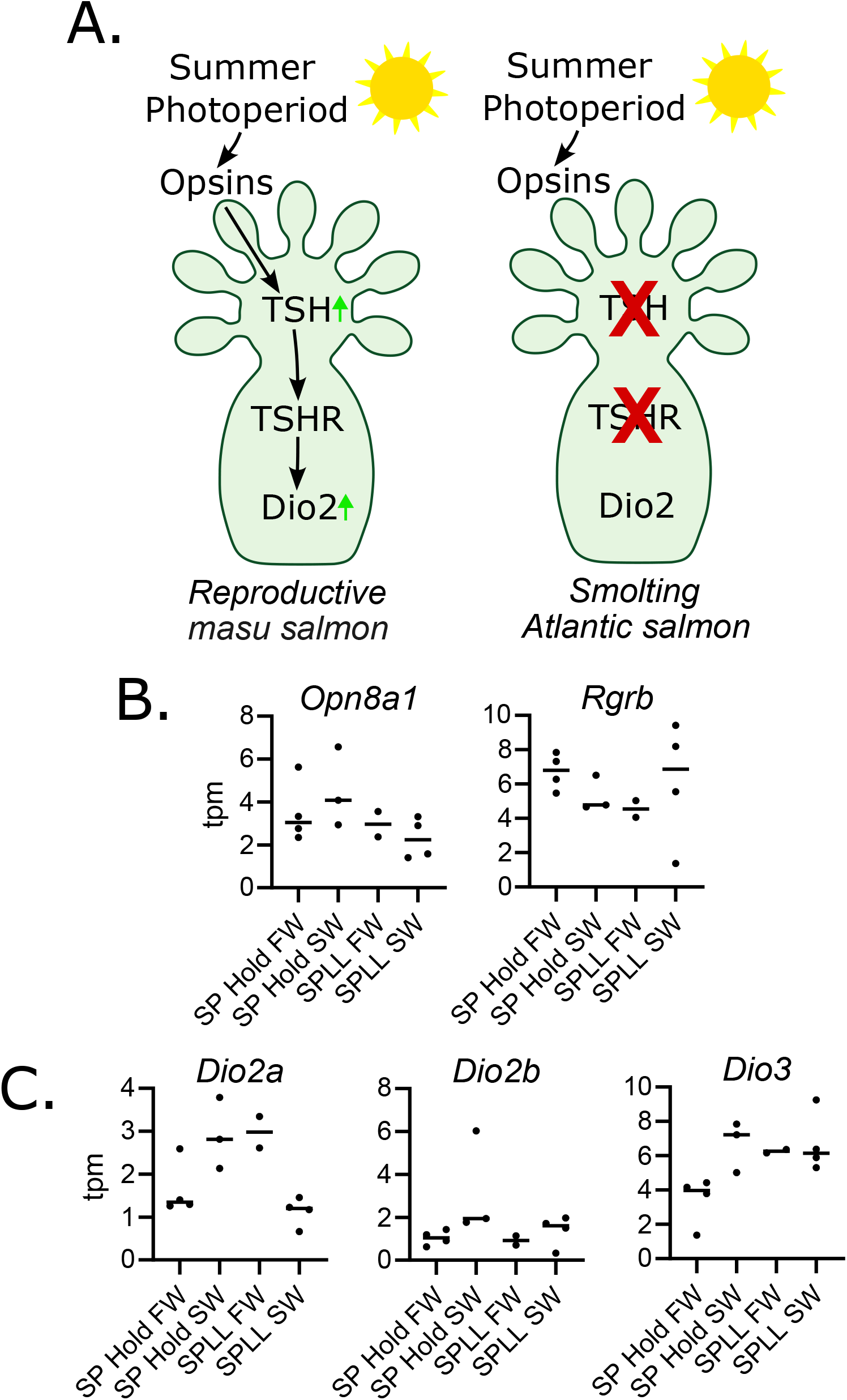
TSH-Dio pathway lacks key elements in the Atlantic salmon saccus vasculosus during parr-smolt transformation (A) schematic showing the pathway from light to increased Dio2 expression identified in precocious male masu salmon^17^ and the comparison to the elements identified in our data of the smoltifing Atlantic salmon. (B) Opsin transcript expression in the Atlantic salmon SV. (C) Deiodinase transcript expression in the Atlantic salmon SV.

## 4. Discussion

To investigate the potential role of the SV during smoltification and seawater transfer in Atlantic salmon, we have characterised the transcriptomic response of the SV to both photoperiod and seawater transfer. Our transcriptomic analysis identified 335 transcripts in the SV whose expression changed in response to photoperiod and / or salinity changes. Apart from genes in cluster 1, which comprise about a fifth of all DEGs detected, the overall message from this analysis is that salinity-dependent changes in SV gene expression are highly dependent on prior photoperiodic history. In some cases, this means a heightened response in smoltified fish (e.g. cluster 3) and in others an attenuated response (e.g. cluster 4). Potentially, these differences stem from differences in the systemic physiological response to SW exposure due to smoltification-related preparatory changes in physiology -for example in the hypo-osmoregulatory capacity of the gill, manifested in differences in plasma osmolality and ionic composition (see **Figure 1**)^61^. Alternatively, they may reflect photoperiodic programming at the level of the SV itself, which then leads to altered responses to ionic, metabolic or hormonal changes upon exposure to SW. Disentangling these scenarios will be difficult, and it is likely that both phenomena contribute to the observed changes in SV responsiveness. In this regard, it is interesting to note that cluster 1 genes (i.e. genes whose expression is increased by SW exposure, independent of photoperiodic history / smoltification status) include a group of circadian clock genes (*Cry1, Tef, Nr1d1* and *Nr1d2)*, which our previous work shows are also induced by seawater transfer in the Atlantic salmon gill ^62^. The similar patterns of upregulation of these transcription factors in two separate tissues suggests that these factors may play a key role in the transcriptional response to seawater entry, distinct from their typical association with circadian clock control (for discussion see ^62^). Potentially this history-independent aspect of the SV response to SW constitutes the primary response pathway, which then triggers the history-dependent downstream changes based on epigenetic differences transcription factor binding site access to other regions of the genome.

While work in the masu salmon has led to the suggestion that the SV may directly integrate photoperiodic information leading to coordination of physiological changes, we have been unable to detect expression of the key ingredients of this putative pathway in the SV transcriptome of the Atlantic salmon. Specifically, while Nakane et al reported expression of the opsins *Rh1, Sws1, Lws* and *Opn4* in SVs of underyearling precocious male Masu salmon, no transcript expression for any of these genes was observed in our dataset. Nevertheless, our detection of *Opn8a1* and *Rgrb* transcripts means that direct light sensitivity in the Atlantic salmon SV remains a possibility. Furthermore, levels of expression of *Tshβa, Tshβb*, the glycoprotein alpha subunit genes *Glha1* and *Gpha2*, and of the thyrotropin receptor (*Tshr*) were all below the detection limit of our RNA seq experiments. While we could detect type 2 and type 3 deiodinase gene expression (*Dio2a, Dio2b & Dio3*) in the SV, no effect of either photoperiodic history or salinity change was observed. Overall, we are forced to conclude that photoperiodic influences on the SV are not tissue-intrinsic responses through an opsin-TSH-DIO cascade, and we therefore favour a model in which smolification-related changes in SV function occur as secondary response to neuroendocrine responses to photoperiod mediated by the hypothalamo-pituitary axis.

While Nakane and colleagues have proposed that the SV is involved in seasonal reproductive regulation in masu salmon, there is no obvious link between the changes in the SV transcriptome observed in the present study and control of the gonadal axis in Atlantic salmon. Rather, several important aspects of our data are consistent with earlier work linking the SV to the regulation of CSF composition. The high contribution (>6 %) of transcripts for the ependymins, *Epn1/2* to the overall transcriptomic signature is of particular interest. Ependymins are the dominant protein component of the CSF in teleosts, and their solubility is calcium-dependent ^63,64^. Previous work highlights the high density of Ca^2+^ containing vesicles in the CSF-contacting globules of the SV coronet cells. These vesicles are thought to be physiologically controlled reservoirs that maintain calcium homeostasis in the cerebrospinal fluid (CSF) ^20–23^. Our results suggest a revision of this hypothesis: the SV is a circumventricular organ responsible for high levels of secretion of ependymin to maintain CSF composition, and the vesicular packaging and solubility of ependymin depend on high levels of calcium in the secretory vesicles of the coronet cells. The extensive innervation of the SV by hypothalamic neuronal projections and by CSF-contacting dopaminergic neurons ^24,25^, as well as the extensive transcriptomic responses of the SV to SW exposure would then reflect the need for precise control of ependymin secretion during the transition to a new aquatic environment.

Taken together, our data shows that in the Atlantic salmon, the SV transcriptome responds to SW exposure in a manner that depends on photoperiodic history and smoltification status. Significantly, most of the key elements of the opsin-TSH-DIO seasonal trigger pathway (eg. *TSHb* subunits, *Glha1, Gpha2* and *TSHR*) are not expressed in the SV, suggesting that in contrast to the proposed role of the SV in seasonal reproduction of masu salmon ^17^, the Atlantic salmon SV does not play a coordinating role in smoltification. Available evidence is consistent with a role for the SV in regulation of CSF composition during transition between freshwater and marine environments.

## 5. Acknowledgements

We thank the diligent care to the fish provided by the staff at Havsbruksstasjonen in Kårvik.

## References

1. Hazlerigg, D. G., Appenroth, D., Tomotani, B. M., West, A. C. & Wood, S. H. Biological timekeeping in polar environments: lessons from terrestrial vertebrates. Journal of Experimental Biology 226, (2023).

2. Mobley, K. B. et al. Maturation in Atlantic Salmon (Salmo Salar, Salmonidae): A Synthesis of Ecological, Genetic, and Molecular Processes. Reviews in Fish Biology and Fisheries vol. 31 (2021).

3. West, A. C., Hazlerigg, D. G. & Grenier, G. Calendar timing in Teleost Fish. In Neuroendocrine clocks and calendars 143–162 (Masterclass in Neuroendocrinology, 2020).

4. West, A. C. et al. Immunologic Profiling of the Atlantic Salmon Gill by Single Nuclei Transcriptomics. Front Immunol 12, 1–11 (2021).

5. Nuñez-Ortiz, N. et al. Atlantic salmon post-smolts adapted for a longer time to seawater develop an effective humoral and cellular immune response against Salmonid alphavirus. Fish Shellfish Immunol 82, 579–590 (2018).

6. Johansson, L. H., Timmerhaus, G., Afanasyev, S., Jørgensen, S. M. & Krasnov, A. Smoltification and seawater transfer of Atlantic salmon (Salmo salar L.) is associated with systemic repression of the immune transcriptome. Fish Shellfish Immunol 58, 33–41 (2016).

7. Stefansson, S. O., Björnsson, B. T., Ebbesson, L. O. E. & McCormick, S. D. Smoltification. In Fish Larval Physiology 639–681 (2008).

8. Nakao, N. et al. Thyrotrophin in the pars tuberalis triggers photoperiodic response. Nature 452, 317–322 (2008).

9. Hanon, E. A. et al. Ancestral TSH Mechanism Signals Summer in a Photoperiodic Mammal. Current Biology 18, 1147–1152 (2008).

10. Masumoto, K. et al. Acute Induction of Eya3 by Late-Night Light Stimulation Triggers TSHβ Expression in Photoperiodism. Curr Biol 20, 2199–2206 (2010).

11. West, A. C. & Wood, S. H. Seasonal physiology: making the future a thing of the past. Curr Opin Physiol 5, 1–8 (2018).

12. Takashi Yoshimura et al. Light-induced hormone conversion of T4 to T3 regulates photoperiodic response of gonads in birds. Nature 426, 178–181 (2003).

13. Nakao, N. et al. Thyrotrophin in the pars tuberalis triggers photoperiodic response. Nature 452, 317–322 (2008).

14. Hanon, E. A. et al. Ancestral TSH Mechanism Signals Summer in a Photoperiodic Mammal. Current Biology 18, 1147–1152 (2008).

15. Masumoto, K. et al. Acute Induction of Eya3 by Late-Night Light Stimulation Triggers TSHβ Expression in Photoperiodism. Curr Biol 20, 2199–2206 (2010).

16. Trudeau, V. L. & Somoza, G. M. Multimodal hypothalamo-hypophysial communication in the vertebrates. Gen Comp Endocrinol 293, 113475 (2020).

17. Nakane, Y. et al. The saccus vasculosus of fish is a sensor of seasonal changes in day length. Nat Commun 4, (2013).

18. Irachi, S. et al. Photoperiodic regulation of pituitary thyroid-stimulating hormone and brain deiodinase in Atlantic salmon. Mol Cell Endocrinol 519, (2021).

19. Fleming, M. S. et al. Functional divergence of thyrotropin beta-subunit paralogs gives new insights into salmon smoltification metamorphosis. Sci Rep 9, 1–15 (2019).

20. Cid, P., Doldan, M. & de Miguel Villegas, E. Morphogenesis of the saccus vasculosus of turbot Scophthalmus maximum: assessment of cell proliferation and distribution of parvalbumin and calretinin during ontogeny. J Fish Biol 87, 17–27 (2015).

21. Graña, P., Huesa, G., Anadón, R. & Yáñez, J. Immunohistochemical study of the distribution of calcium binding proteins in the brain of a chondrostean (acipenser baeri). Journal of Comparative Neurology 520, 2086–2122 (2012).

22. Jansen, W. F., Burger, E. H. & Zandbergen, M. A. Subcellular localization of calcium in the coronet cells and tanycytes of the saccus vasculosus of the rainbow trout, Salmo gairdneri Richardson. Cell Tissue Res 224, 169–180 (1982).

23. Jansen, W., Flight, W. & Zandbergen, M. Fine structural localization of adenosine triphosphate activities in the saccus vasculosus of the rainbow trout, Salmo gairdneri Richardson. Cell Tissue Res 219, 267–279 (1981).

24. Joy, K. & Sathyanesan, A. A histoenzymological study of the saccus vasculosus of the freshwater teleost, Clarias batrachus (L.). Z Mikrosk Anat Forsch 93, 297–304 (1979).

25. Sueiro, C. et al. New insights on saccus vasculosus evolution: A developmental and immunohistochemical study in elasmobranchs. Brain Behav Evol 70, 187–204 (2007).

26. Tsuneki, K. A Systematic Survey of the Occurrence of the Hypothalamic Saccus Vasculosus in Teleost Fish. Acta Zoologica 73, 67–77 (1992).

27. Köwitsch, A., Zhou, G. & Groth, T. Medical application of glycosaminoglycans: a review. J Tissue Eng Regen Med 12, e23–e41 (2018).

28. Emanuelsson, H. & von Mecklenburg, C. Metabolic activity in the saccus vasculosus of the rainbow trout, Salmo gairdneri (Richardson). Z. Zellforsch 130, 351–361 (1972).

29. Benjamin, M. Ultrastructural Studies on the Coronet Cells of the Saccus vasculosus of the Freshwater Stickleback, Gasterosteus aculeatus form leiurus. Z. Zellforsch 147, 551–565 (1974).

30. Emanuelsson, H. & von Mecklenburg, C. Experimental and structural analysis of the function of saccus vasculosus in rainbow trout, Salmo gairdneri (Richardsen). Cell Tiss. Res. 148, 27–44 (1974).

31. Martin, M. Cutadapt removes adapter sequences from high throughput sequencing reads. EMBnet Journal 17, 10–12 (2011).

32. Patro, R., Duggal, G., Love, M. I., Irizarry, R. A. & Kingsford, C. Salmon: fast and bias-aware quantification of transcript expression using dual-phase inference. Nat Methods 14, 417–419 (2017).

33. Robinson, M. D., McCarthy, D. J. & Smyth, G. K. edgeR: A Bioconductor package for differential expression analysis of digital gene expression data. Bioinformatics 26, 139–140 (2009).

34. Smedley, D. et al. BioMart - Biological queries made easy. BMC Genomics 10, 1–12 (2009).

35. Eden, E., Navon, R., Steinfeld, I., Lipson, D. & Yakhini, Z. GOrilla: A tool for discovery and visualization of enriched GO terms in ranked gene lists. BMC Bioinformatics 10, 1–7 (2009).

36. Mulugeta, T. D. et al. SalMotifDB: a tool for analyzing putative transcription factor binding sites in salmonid genomes. BMC Genomics 20, 694 (2019).

37. Nilsen, T. O. et al. Differential expression of gill Na+,K+-ATPase α- and β-subunits, Na+,K+,2Cl-cotransporter and CFTR anion channel in juvenile anadromous and landlocked Atlantic salmon Salmo salar. Journal of Experimental Biology 210, 2885–2896 (2007).

38. Mccormick, S. D., Regish, A. M., Christensen, A. K. & Björnsson, B. T. Differential regulation of sodium – potassium pump isoforms during smolt development and seawater exposure of Atlantic salmon. J Exp Biol 216, 1142–1151 (2013).

39. Iversen, M. et al. RNA profiling identifies novel, photoperiodhistory dependent markers associated with enhanced saltwater performance in juvenile Atlantic salmon. PLoS One 15, 1–21 (2020).

40. Hoffmann, W. & Schwarz, H. Ependymins: Meningeal-derived extracellular matrix proteins at the blood-brain barrier. Int Rev Cytol 165, 121–123 (1996).

41. Maeda, R., Shimo, T., Nakane, Y., Nakao, N. & Yoshimura, T. Ontogeny of the saccus vasculosus, a seasonal sensor in fish. Endocrinology 156, 4238–4243 (2015).

42. Schwaller, B. The continuing disappearance of “pure” Ca2+ buffers. Cell. Mol. Life Sci. 275–300 (2009).

43. Nigro, P., Pompilio, G. & Capogrossi, M. Cyclophilin A: a key player for human disease. Cell Death Dis 4, (2013).

44. Lang, K., Schmid, F. X. & Fischer, G. Catalysis of protein folding by prolyl isomerase. Nature 329, 268–270 (1987).

45. Chakrabarti, P. & Khatun, R. Histophysiological and surface ultrastructural studies of the saccus vasculosus of Notopterus chitala (Hamilton). J Microsc Ultrastruct 5, 140 (2017).

46. Grad, I. & Picard, D. The glucocorticoid responses are shaped by molecular chaperones. Molecular and Cellular Endocrinology vol. 275 2–12 Preprint at 10.1016/j.mce.2007.05.018 (2007).

47. Riccardi, C. GILZ as a mediator of the anti-inflammatory effects of glucocorticoids. Frontiers in Endocrinology vol. 6 Preprint at 10.3389/fendo.2015.00170 (2015).

48. Fonjallaz, P., Ossipow, V., Wanner1, G. & Schibler2, U. The Two PAR Leucine Zipper Proteins, TEF and DBP, Display Similar Circadian and Tissue-Specific Expression, but Have Different Target Promoter Preferences. The EMBO Journal vol. 15 https://www.embopress.org (1996).

49. Sigoillot, S. M. et al. The secreted protein C1QL1 and its receptor BAI3 control the synaptic connectivity of excitatory inputs converging on cerebellar purkinje cells. Cell Rep 10, 820–832 (2015).

50. Svenningsson, P. et al. DARPP-32: An Integrator of Neurotransmission. Annual Review of Pharmacology and Toxicology vol. 44 269–296 Preprint at 10.1146/annurev.pharmtox.44.101802.121415 (2004).

51. Zeng, H., Kaul, S., Stoney, S. & Jr, S. Genomic Organization of Human GMEB-1 and Rat GMEB-2: Structural Conservation of Two Multifunctional Proteins. Nucleic Acids Research vol. 28 (2000).

52. Vatine, G. et al. Light directs zebrafish period2 expression via conserved D and E boxes. PLoS Biol 7, (2009).

53. West, A. C. et al. Diversified regulation of circadian clock gene expression following whole genome duplication. PLoS Genet 1–21 (2020) doi:10.1371/journal.pgen.1009097.

54. Wood, S. H. et al. Binary switching of calender cells in the pituitary defines the phase of the circannual cycle in mammals. Current Biology 25, (2015).

55. Kao, S. H., Wu, H. T. & Wu, K. J. Ubiquitination by HUWE1 in tumorigenesis and beyond. Journal of Biomedical Science vol. 25 Preprint at 10.1186/s12929-018-0470-0 (2018).

56. Madison, B.B. Srebp2: A master regulator of sterol and fatty acid synthesis1. Journal of Lipid Research vol. 57 333–335 Preprint at 10.1194/jlr.C066712 (2016).

57. Hara, S. et al. Prostaglandin E synthases: Understanding their pathophysiological roles through mouse genetic models. Biochimie vol. 92 651–659 Preprint at 10.1016/j.biochi.2010.02.007 (2010).

58. Eilertsen, M. et al. Photoreception and transcriptomic response to light during early development of a teleost with a life cycle tightly controlled by seasonal changes in photoperiod. PLoS Genet 18, 1–28 (2022).

59. Eilertsen, M. et al. An EvoDevo Study of Salmonid Visual Opsin Dynamics and Photopigment Spectral Sensitivity. Front Neuroanat 16, 1–15 (2022).

60. Fleming, M. S. et al. Functional divergence of thyrotropin beta-subunit paralogs gives new insights into salmon smoltification metamorphosis. Sci Rep 9, 1–15 (2019).

61. Hoar, W. S. THE PHYSIOLOGY OF SMOLTING SALMONIDS. (1988).

62. West, A. C. et al. Diversified regulation of circadian clock gene expression following whole genome duplication. PLoS Genet 1–21 (2020) doi:10.1371/journal.pgen.1009097.

63. Shashoua, VE. Ependymin, a Brain Extracellular Glycoprotein, and CNS Plasticity. Ann N Y Acad Sci 627, 94–114 (1991).

64. Hoffmann, W. & Schwarz, H. Ependymins: Meningeal-derived extracellular matrix proteins at the blood-brain barrier. Int Rev Cytol 165, 121–123 (1996).

65. Iversen, M. et al. RNA profiling identifies novel, photoperiodhistory dependent markers associated with enhanced saltwater performance in juvenile Atlantic salmon. PLoS One 15, 1–21 (2020).

